# Generation of Super-resolution Images from Barcode-based Spatial Transcriptomics Using Deep Image Prior

**DOI:** 10.1101/2023.06.26.546529

**Authors:** Jeongbin Park, Seungho Cook, Dongjoo Lee, Jinyeong Choi, Seongjin Yoo, Hyung-Jun Im, Daeseung Lee, Hongyoon Choi

## Abstract

Spatial transcriptomics (ST) has revolutionized the field of biology by providing a powerful tool for analyzing gene expression *in situ*. However, current ST methods, particularly barcode-based methods, have limitations in reconstructing high-resolution images from barcodes sparsely distributed in slides. Here, we present SuperST, a novel algorithm that enables the reconstruction of dense matrices from low-resolution ST libraries. SuperST based on deep image prior reconstructs spatial gene expression patterns as image matrices. SuperST allows gene expression mapping to better reflect immunofluorescence (IF) images. Compared with previous methods, SuperST generated output images that more closely resembled IF images for given gene expression maps. Additionally, SuperST overcomes the limitations inherent in IF images, highlighting its potential applications in the realm of spatial biology. By providing a more detailed understanding of gene expression *in situ*, SuperST has the potential to contribute to comprehensively understanding biology from various tissues.

## Introduction

Spatially resolved transcriptomics (ST) becomes an emerging technology in biology, facilitating the discovery of new drugs and pathology. There are currently two types of ST methods: spot-based ST and image-based ST^1^. Spot-based ST, such as 10x Visium^2^, differs from image-based ST^1^, by providing whole gene-level expression data and typically involves thousands of spots. However, there are tens of cells in a spot, limiting its resolution and making it difficult to interpret underlying biology and cellular level interpretation. Specifically, barcode-based ST has limitations, as it can only collect data from specified positions and struggles to interpret empty spaces. Interpreting the positions of barcodes is difficult, making it challenging to treat the barcodes as a dense image matrix and apply image processing tools such as image registration and integration^3, 4^. Additionally, RNA diffusion in tissues during experiments limits the resolution of even image-based ST platforms. The reported velocities of moving RNAs or RNA-protein complexes were ranged from 0.65 to 1.5 μm/s^5–8^ depending on tissues, conditions, and molecules of interest.

To date, several algorithms have been developed to improve the resolution of ST. BayesSpace used the Bayesian approach to predict sub-spot gene expression, considering the structure of neighboring spots^9^. The spatially-aware dimension reduction method (SpatialPCA) extracted a low-dimensional representation using probabilistic principal component analysis (PCA) and predicted gene expression in unmeasured spots in a spatially-aware imputation manner^10^. Deep spatial data fusion, also known as XFuse, was a histology-dependent super-resolution image generation algorithm using a deep-generative model for Visium spatial transcriptomics^11^. A recent algorithm of tumor edge structure and lymphocyte multi-level annotation (TESLA) involves determining neighborhood relationships from H&E images and imputing super-pixels to generate super-resolution gene expression images^12^. There have also been various attempts to assume gene expression in unknown regions by introducing well-known probabilistic models, such as Gaussian distribution, negative binomial distribution, and Poisson distribution.

Despite the existence of several methods for improving ST resolution, the reconstruction of dense image matrices from spot-based data remains a challenge. However, by predicting gene expression data as images, it is possible to apply various image processing methods (*e.g.*, segmentation^13^, registration, contouring, and transformation) to utilize spatial transcriptomics data as well as obtain high-resolution data. This approach holds great potential for advancing the field of ST and expanding our understanding of gene expression *in situ*. In this regard, we proposed and validated the algorithm, SuperST, for creating Super-resolution images by applying deep image prior from low-resolution ST libraries.

## Results

### DIP produced image matrices of gene expression from ST data

SuperST was based on an exemplary deep image prior (DIP) algorithm^14^ (**Figure. 1a**). Briefly, DIP is a deep learning approach that reconstructs high-quality images from low-quality inputs, such as noisy or blurred images, without the need for pre-labeled training data. It uses an untrained neural network, for here, U-Net^15^, to constrain the image reconstruction process, resulting in high-resolution images. For DIP, there was two modules in the algorithm: down-sampling (colored in orange) and up-sampling (colored in blue) in **Figure. 1a**. The input for this process was the histology image, and the output was generated through U-Net. Specifically, we designated the matrix size of gene expression map, and set to 256×256 matrix for each gene. We resized the histology image, the input for U-Net, to 256×256 and fed it into the U-Net to predict a set of gene expression matrices of specific genes. We then applied a convolution term of Gaussian smoothing to account for the diffusion of mRNA. This term considered the diffusion of mRNA expression in the periphery for each position, which was counted as gene expression for each position of the spot. The output 256×256 image was then masked only with pixels corresponding to the central position of each spot, and a loss function was created with the difference between the value of each masked pixel and the corresponding gene expression value for each spot. To minimize this loss function, the model was iteratively trained. The output became the gene expression value of the dense matrix immediately before masking. The example of the human breast cancer sample^16^ was presented in **Figure. 1b**.

**Figure 1.**
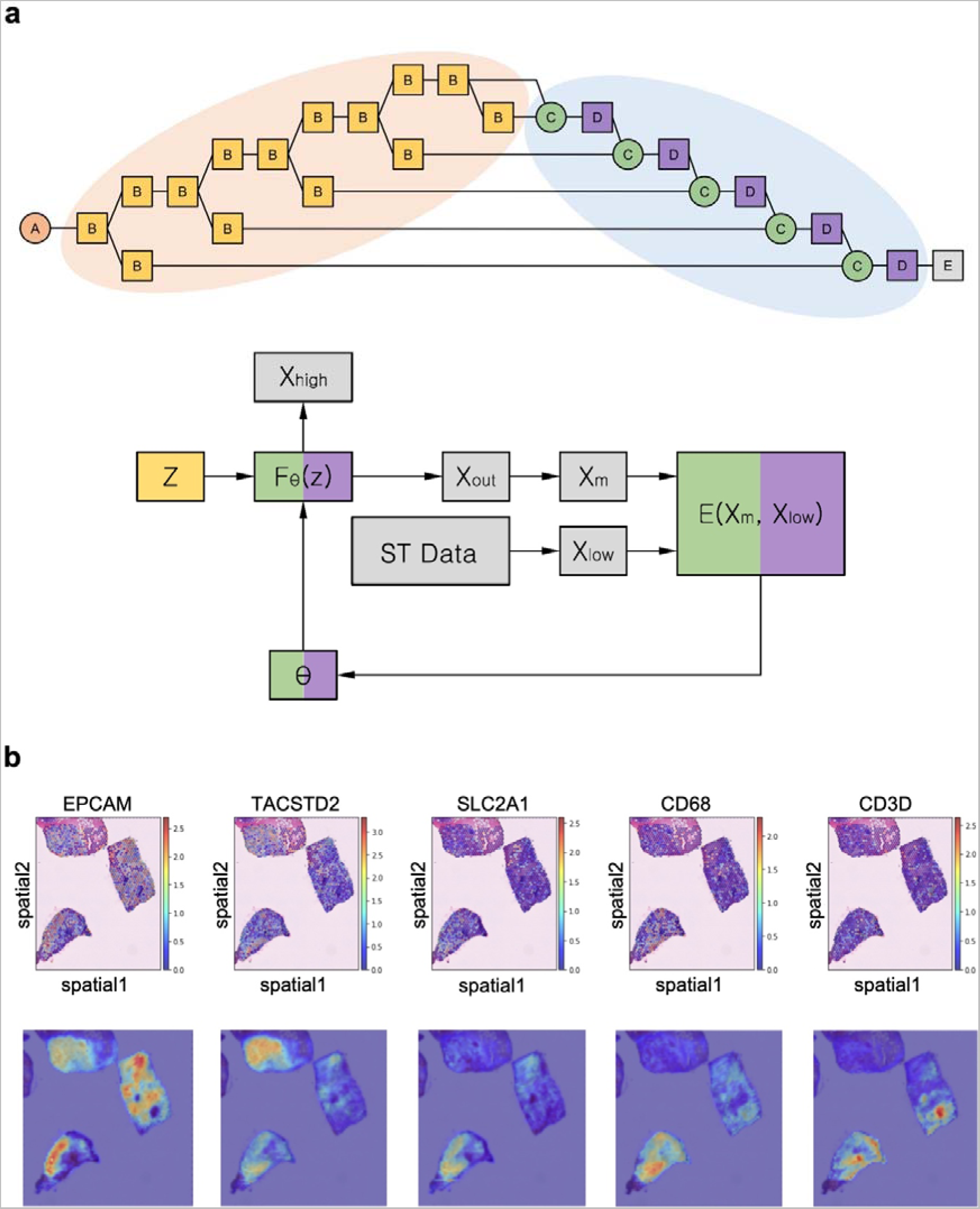
The design of SuperST. **a**, The schematic representation of SuperST. Here, A, B, C, D, and E represent ‘*input_1*’ representing an input H&E image, conceptual down-sampling unit, ‘*concatenate*’, conceptual up-sampling unit, and the output of U-Net. The algorithm runs with F_θ_(z) → X_out_ by ‘*num_iter*’ times of execution at first for updating U-Net and F_θ_(z) → X_high_ once for predicting high-resolution image. F denotes a conceptual function that links the CNN architectures (B) to the output (E). **b**, The comparison of conventional spatial feature plots and high-resolution images made by SuperST from a publicly available breast cancer dataset. The pixels in high-resolution images darker than 95 percentiles of each image were not shown.

### Optimal parameters for SuperST

In order to evaluate the performance of SuperST in predicting image matrix of specific gene expression data, we compared the output images obtained from SuperST with immunofluorescence (IF) images. We assessed the correspondence between the output of SuperST generated from a specific gene expression image and the immunofluorescence image obtained from the tissue (**Figure. 2a**). This was achieved by evaluating the pixelwise correlation between the two images. When exploring image similarity metrics including Pearson correlation coefficient (‘*pearson*’) and mutual information (‘*mutual’*), the variation of overall coefficients decreased when the number of iteration (‘*num_iter*’) increased, but the similarity tended to decrease further beyond a certain number of iterations (**Supplementary Figure. 1**). This could be due to the overfitting of SuperST as can be seen in the case that ‘*num_iter*’ was larger than 1024 (**Supplementary Figure. 2**). However, with a small number of ‘*num_iter*’, the output image did not accurately represent the input gene expression. For example, *S100a9*, a well-known marker gene for neutrophils, was expressed in the inner necrotic core of the 4T1 tumor^17^. However, when ‘*num_iter*’ was less than 128, the output images did not show the predominant *S100a9* expression in the core region other than the conventional spatial mapping (**Supplementary Figure. 3**). Therefore, the use of the ‘*num_iter*’ parameter involves a trade-off. If this parameter is set to a high value, it can cause the output images too responsive to the sparse gene expression. On the other hand, setting ‘*num_iter*’ to a low value can lead to an underestimation of gene expression information. However, when ‘*num_iter*’ is set within an appropriate range, the performance of SuperST is optimized and relatively constant. Therefore, the semi-parameter independence approach can be helpful in selecting the appropriate ‘*num_iter*’ value for the specific application.

**Figure 2.**
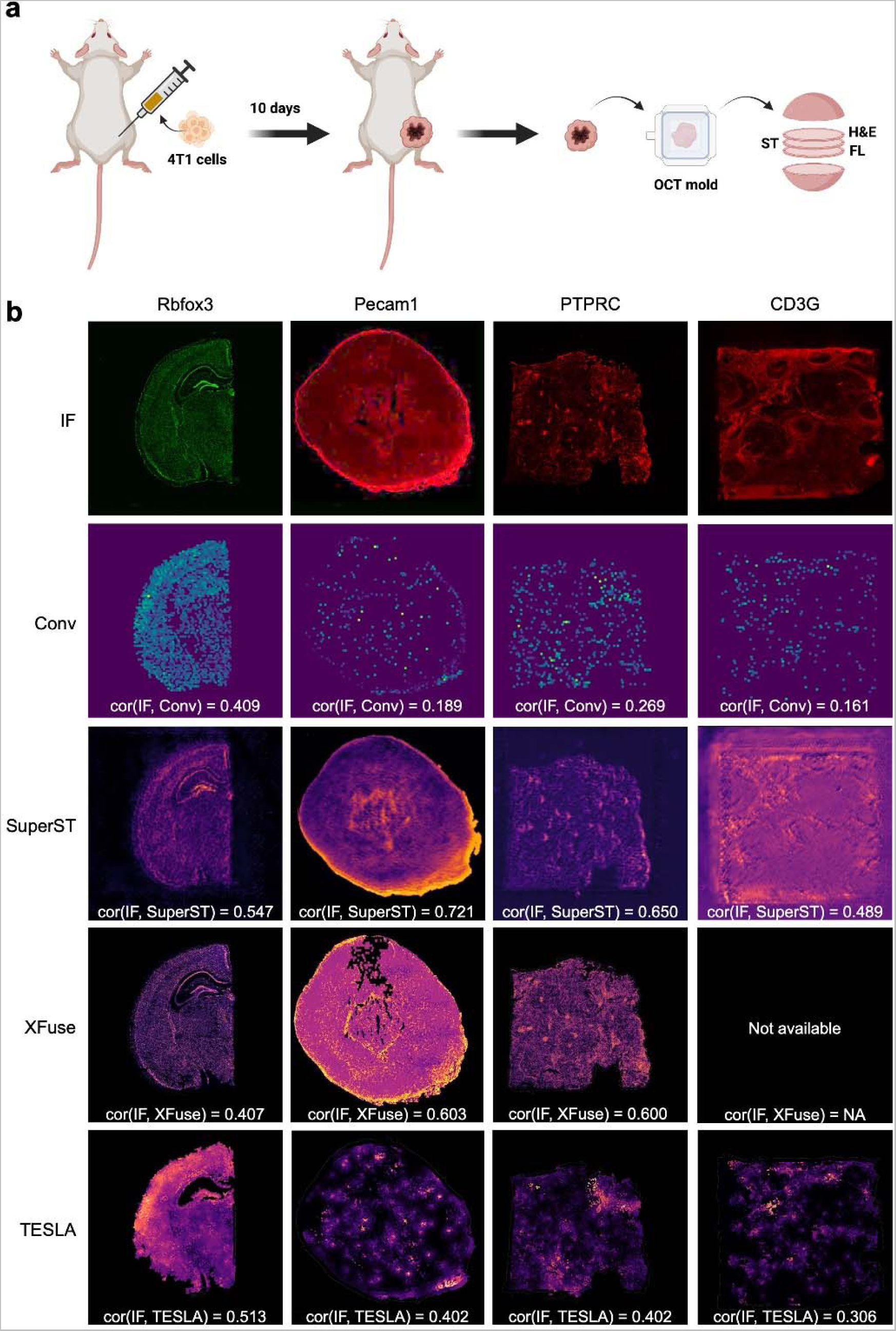
Comparison between conventional visualization, SuperST, XFuse, and TESLA with the IF image. **a**, The acquisition of Visium ST library for ‘*Mouse 4T1*’ with a recommended tissue preparation protocol, followed by the primary and the fluorescent secondary antibody treatment. H&E, ST, and FL represented the H&E-stained tissue, the spatial transcriptomics library, and the fluorescent tissue. **b**, Each Pearson correlation coefficient between IF and each super-resolution algorithm (SuperST, XFuse, and TESLA) output was represented for ‘*Mouse Brain*’ (*Rbfox3*), ‘*Mouse 4T1*’ (*Pecam1*), ‘*Human Ovarian*’ (*PTPRC*), and ‘*Human Ductal*’ (*CD3G*). ‘Conv’ indicating the conventional visualization of ST data (negative control) was also compared with IF. Super-resolution algorithms outperformed the conventional visualization in most cases. Also, the correlation coefficient was found to be the highest in SuperST compared with the other algorithms.

### Comparison with previous methods

As previous methods to provide relatively high-resolution image matrix from ST data, XFuse and TESLA were performed for four different datasets, along with SuperST and conventional visualization of ST data (**Figure. 2b**). It was noteworthy that the output images of the super-resolution algorithms better reflected the gene expression profiles than sparse Visium expression images, which poorly exhibited continuous actual gene expression. SuperST showed the highest Pearson correlation coefficient between the output images and the IF images. XFuse and TESLA exhibited inappropriately contoured outputs in several use cases (**Figure. 2b**, *Pecam1* for XFuse and *Rbfox3* for TESLA). Also, there was no available output for certain genes (*e.g.*, *CD3G*) when running XFuse. Moreover, discrepancies were identified between the output images of XFuse or TESLA and the IF images. For example, in the ‘*Mouse 4T1*’ sample, the inner necrotic section and the outer stromal compartment in the anti-Pecam1 antibody IF image were more accurately depicted in SuperST compared with XFuse and TESLA. Also, of all the methods, TESLA generates results that are less similar to the IF images, although it still surpasses the conventional visualization in performance for all the cases.

## Discussion

The SuperST algorithm proposed in this study aimed to address the challenge of reconstructing dense image matrices from spot-based ST data, by predicting gene expression data as images using the DIP algorithm. The study demonstrated that DIP produced high-resolution image matrices of gene expression from low-resolution ST data. The study evaluated the performance of SuperST by comparing the output images obtained from SuperST with IF images and assessing the correspondence between the two types of images using image similarity metrics. The results showed that SuperST generated output images that were highly correlated with IF images.

The proposed algorithm has several significant implications for the field of spatial transcriptomics. It provides a way to overcome the limitations of barcode-based ST, which can only collect data from specified positions and struggles to interpret empty spaces. The use of DIP in SuperST allows for the reconstruction of high-resolution images from low-resolution inputs, which could potentially revolutionize the field of ST by facilitating the discovery of new targets and pathophysiology of various diseases.

While interpolation assumes a specific spatial distribution for gene expression, such as Gaussian distribution, negative bionomical distribution, and Poisson distribution, super-resolution algorithms including SuperST did not rely on the assumptions. SuperST can generate varying output images in a probabilistic way, and it can also provide clues for optimal parameters through semi-parameter independence. After searching for the best parameters, we conducted a comparison between the SuperST results and IF images to examine the molecular expression in the tissue. It is noteworthy that IF detects proteins that are distinct from mRNA transcripts of ST^18, 19^. Nonetheless, our findings indicated that SuperST exhibited a stronger correlation with the spatial patterns observed in IF, surpassing other methods (*i.e.*, XFuse and TESLA).

Additionally, we observed that SuperST effectively eliminated unreliable signals that were present in the IF images. We compared IF images and SuperST outputs (**Supplementary Figure. 4**). Fluorescence images, including IF images, were prone to show autofluorescence, which reduces the reliability of the images. Even in the absence of fluorescent materials, autofluorescence could be observed, especially at the edges of tissue images (**Supplementary Figure. 4**, white arrow). It is remarkable that SuperST successfully differentiated between autofluorescence and genuine gene expression by effectively representing gene expression information. However, additional assessment was still required to determine if the edge signal was indeed autofluorescence. In addition, the antibody-staining technique used in IF images were also affected by histological conditions, such as the nonspecific binding of antibodies^20^. However, SuperST reduced the impact of histological conditions on the IF images, as evidenced by the absence of scarred tissue and duct-specific patterns in the SuperST outputs compared with the IF images (**Supplementary Figure. 4**, yellow arrow). In this regard, SuperST effectively served as a regularizer, striking a balance between gene expression and histology images, thereby generating more precise molecular expression patterns.

The study also has some limitations that need to be considered. The SuperST algorithm was evaluated using limited samples, which may hardly support the generalizability of the results to whole gene expression of other tissues. Additionally, the proposed algorithm requires key parameters for DIP, which need to be optimized for generalization. It was discovered that the number of iteration (‘*num_iter*’) is crucial and needs to fall within a specific range. However, it also needs to be customized according to the characteristics of the data provided. Also, this study focuses solely on ‘*num_iter*’ for optimal SuperST performance, but other factors like image size, kernel size, and sigma value should be tuned for better performance. Despite these limitations, the SuperST algorithm has great potential for advancing the field of ST and expanding our understanding of gene expression *in situ*.

In conclusion, our results demonstrate that SuperST could generate dense image matrices with high-resolution information from barcode-based ST data. SuperST outperformed existing methods by reflecting gene expression and histological information properly. In terms of future perspectives, SuperST has the potential to be applied in a wide range of research fields that analyze ST leveraging imaging analysis workflows. For example, it could be used in various image processing algorithms for analyzing ST data by changing the sparsely distributed matrix data into high-resolution image data. Furthermore, the image data generated by SuperST could be easily integrated with other spatial omics technologies, such as proteomics and metabolomics, which could lead to a more comprehensive understanding of biological systems. Overall, SuperST represents a significant advancement in the field of ST and has the potential to transform our understanding of complex biological systems.

## Methods

### Datasets

A public breast cancer dataset was used to compare the results with and without SuperST^16^. Three public datasets were additionally utilized for the validation of SuperST. One dataset (‘*Mouse Brain*’) was ‘Adult Mouse Brain Section 2 (Coronal)’ data publicly available from 10x Genomics^21^. The tissue in the dataset was treated with anti-GFAP antibody (glial fibrillary acidic protein, red) and anti-NeuN (mature neuron marker, green) antibody along with DAPI staining (blue). Also, two datasets provided by 10x Genomics including the human ovarian cancer sample (‘*Human Ovarian*’; CD45: Cy5 - red)^22^ and the human invasive ductal carcinoma sample (‘*Human Ductal*’; CD3: Alexa Fluor 647 - red)^23^ were used.

Moreover, we generated another ST dataset (‘*Mouse 4T1*’) from a 4T1-bearing female Balb/c mouse which was prepared by injecting 4T1 tumor cells into the right thigh region of the mouse. The tumor sample was collected and frozen 10 days after injection. Three tissue slices were acquired from the tumor tissue. One slice was treated with anti-PECAM1 primary antibody and red fluorescent anti-IgG secondary antibody. The immunofluorescence (IF) image was then obtained by observing it with confocal microscope (STELLARIS 5, Leica microsystems). Another slice was used for the H&E staining. The third slice was treated with methanol, frozen, cryo-sectioned, and used to generate a ST library. The preparation of the ST library followed a recommended Visium protocol of 10x Genomics.

### Construction of SuperST

SuperST generates a high-resolution image X_high_ from a low-resolution expression matrix X_low_ by employing U-Net to realize deep image prior (DIP) (**Figure. 1a**). The equation (1) illustrates the mechanism of SuperST.

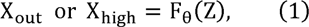

Here, Z ∈ □ ^C’^ ^×^ ^H’^ ^×^ ^W’^ is C’ numbers of CNN architectures of height H’ and width W’ learned from X_0_.

X_out_ or X_high_ ∈ □ ^3^ ^×^ ^H^ ^×^ ^W^ = □ ^3^ ^×^ ^256^ ^×^ ^256^ is the image derived from Z. θ is a network parameter and F is a function made up of deep learning network. X_0_ is allocated to X_low_ in the original DIP, but we used a histology image instead with the following reasons. At first, a histology image has a lot of information so that it can help to predict the high-resolution expression image of a single gene. Second, every ST Visium library has its corresponding histology image, so it is desirable to actively utilize the image.

After generating an output high-resolution image X_out_ from F, it is converted to low-resolution matrix X_m_ to be compared with the low-resolution ST gene expression matrix of a gene, X_low_. In the original DIP, θ is allocated to the determinator θ* for the minimum cost function of E between X_m_ and X_low_ in the equation (2).

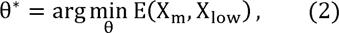

When the number of iterations (‘*num_iter*’) is low, the resultant images did not fully reflect the gene expression pattern. When ‘*num_iter*’ is high, however, the resultant images are too responsive to the sparse gene expression. In an appropriate range of ‘*num_iter*’, the performance of SuperST is optimized and relatively constant. Thus, we chose ‘*num_iter*’ within the interval based on the semi-parameter independence.

### Comparison between SuperST and IF

When performing SuperST for ‘*Mouse Brain*’ (Rbfox3), ‘*Human Ovarian*’ (PTPRC), and ‘*Human Ductal*’ (CD3G), the input images were derived by subtracting the fluorescent antibody channel from the given 3-channel images. Also, for ‘*Mouse 4T1*’ (Pecam1), the H&E staining image was used as the input. With the ST samples, the following image similarity metrics were calculated: Pearson correlation coefficient (‘*pearson*’) and mutual information (‘*mutual*’). When calculating each metric, it was calculated 20 times for each *‘num_iter*’ and all the required parameters were set to the default. Meanwhile, qualitative comparisons were carried out between SuperST and IF with the ST samples.

### Comparison between SuperST and existing methods

The conventional visiualization methods of Visium were performed by enlarging the spot-wise expression in a square form. The inputs for XFuse and TESLA were the same as those for SuperST. XFuse was prepared in a default setting according to the literature^11^. Also, among the XFuse outputs – ‘stdv’, ‘mean’, and ‘invcv+’ images, the ‘invcv+’ image was selected as the representative output of XFuse. In addition, TESLA was run with ‘*apertureSize*’ of 5, ‘*res*’ of 3, and the first type of contouring algorithm of three. Then, we proceeded to compare various super-resolution algorithms using the four datasets: ‘*Mouse Brain*’, ‘*Mouse 4T1*’, ‘*Human Ovarian*’, and ‘*Human Ductal*’. After creating the output images for each dataset, we calculated the Pearson correlation coefficient to measure the relationship between each IF image and its corresponding output image. This was done individually for each algorithm and sample.

### Data and code availability

The generated ST dataset for this study, ‘*Mouse 4T1*’, is available upon a reasonable request. Otherwise, all the other datasets used in this study are publicly available. Additionally, the necessary codes and requirements can be accessed in our GitHub repository at https://github.com/portrai-io/SuperST.

## Author Contributions

Hongyoon Choi designed this project and conceived the basic code. Jeongbin Park modified and revised it for practical uses. Also, Jeongbin Park explored the application of the algorithm for ST in several use cases. Seungho Cook advised the direction of the study. Dongjoo Lee reviewed the algorithm. Animal experiments were carried out and the ST Visium library was acquired by Jeongbin Park and Jinyeong Choi. Seongjin Yoo built an environment to run the algorithm. The manuscript was written by J.P., H-J.I, D.L., and H.C., and all authors have contributed to the completion of the manuscript.

## Competing Interests

Jeongbin Park, Seungho Cook, Dongjoo Lee, Jinyeong Choi, and Seongjin Yoo are researchers in Portrai, Inc. Hyung-Jun Im, Daeseung Lee, and Hongyoon Choi are the co-founders of Portrai, Inc. The schematic illustration was drawn by BioRender.

## Ethics Declaration

Animal experiments for this study were approved by the Woojung Bio IACUC of the Republic of Korea with the approval code of IACUC2001-003. There were no other ethical issues for this study.

## Figure Legends

**Supplementary Figure 1.**
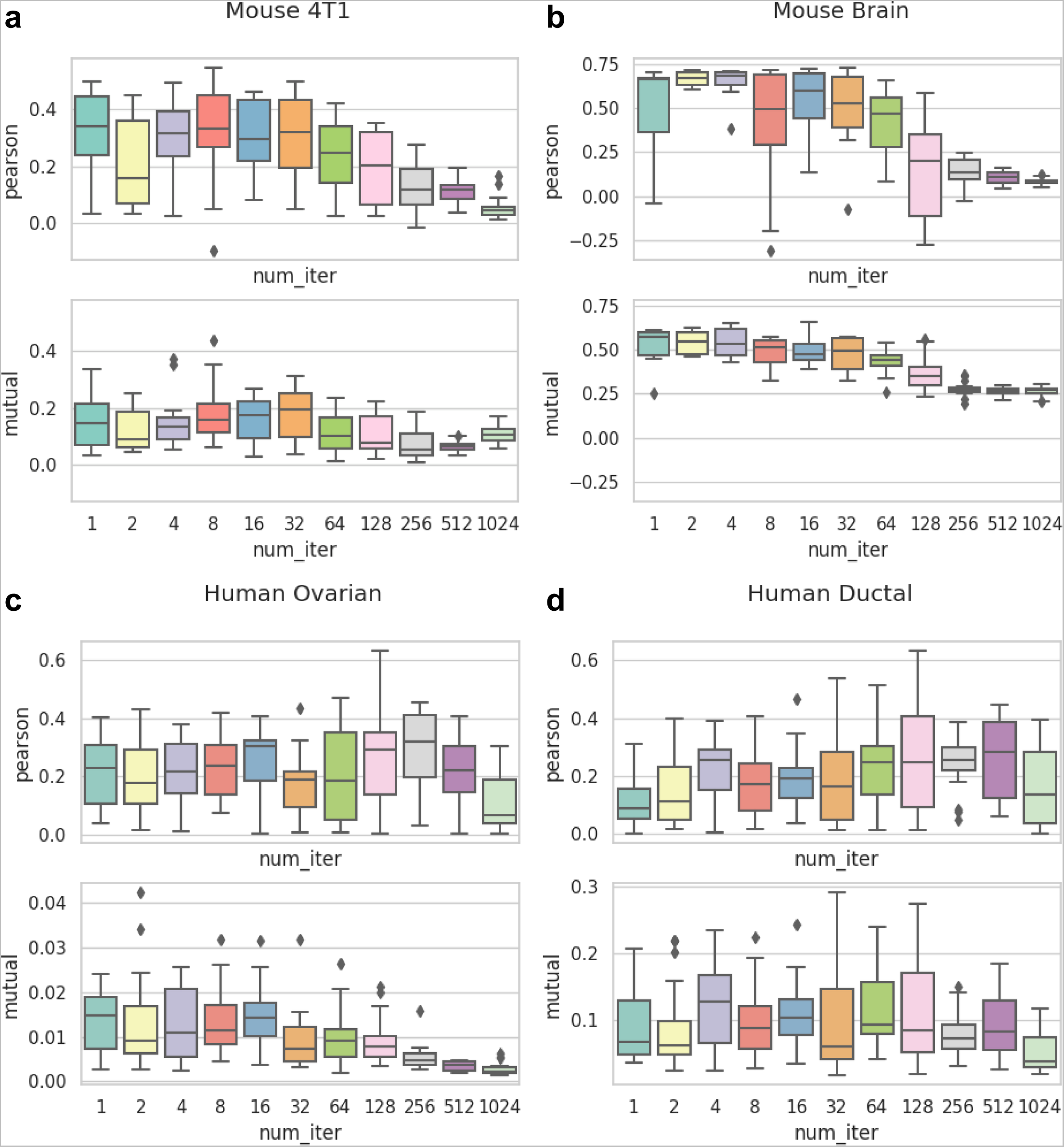

**Supplementary Figure 2.**
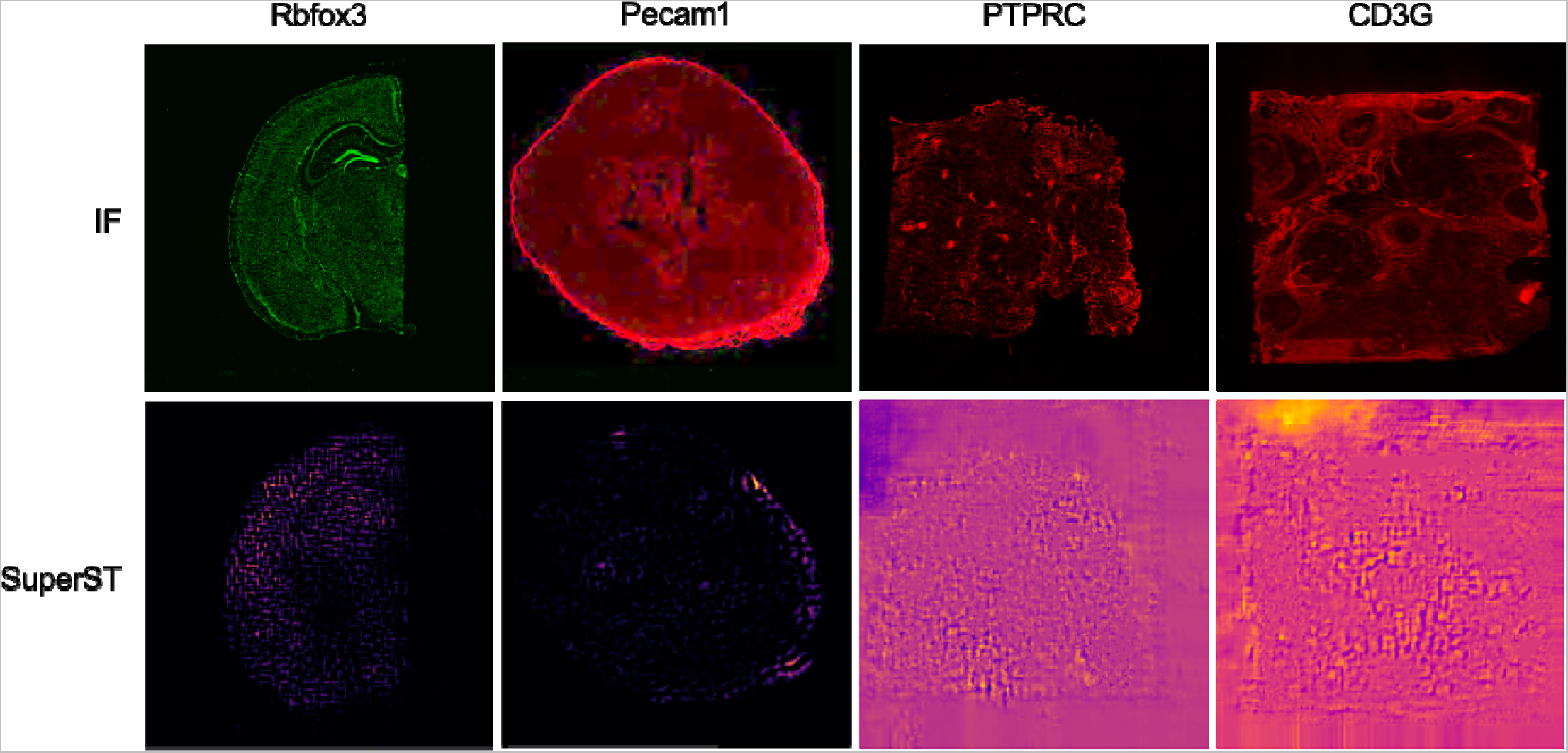

**Supplementary Figure 3.**
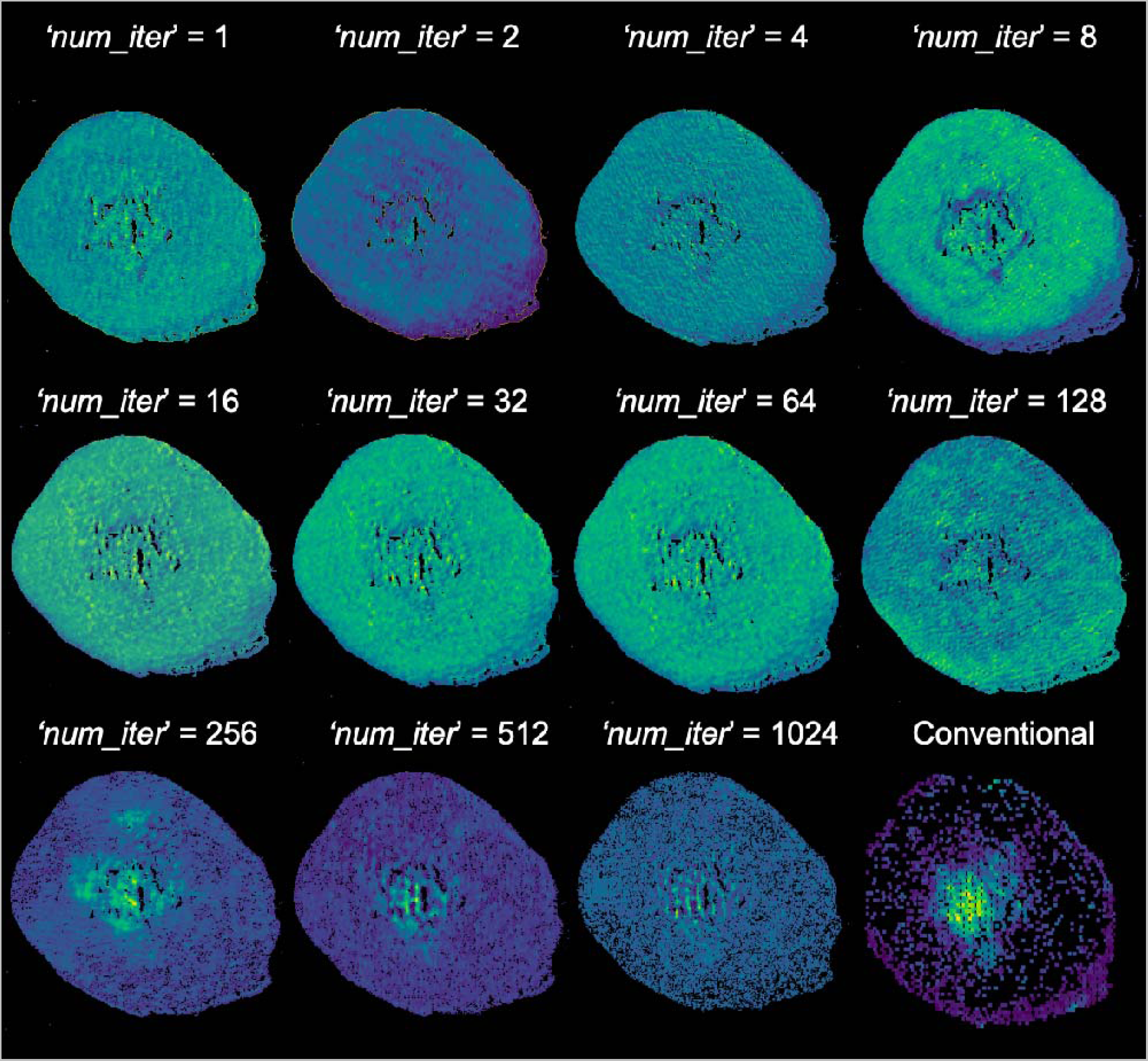

**Supplementary Figure 4.**
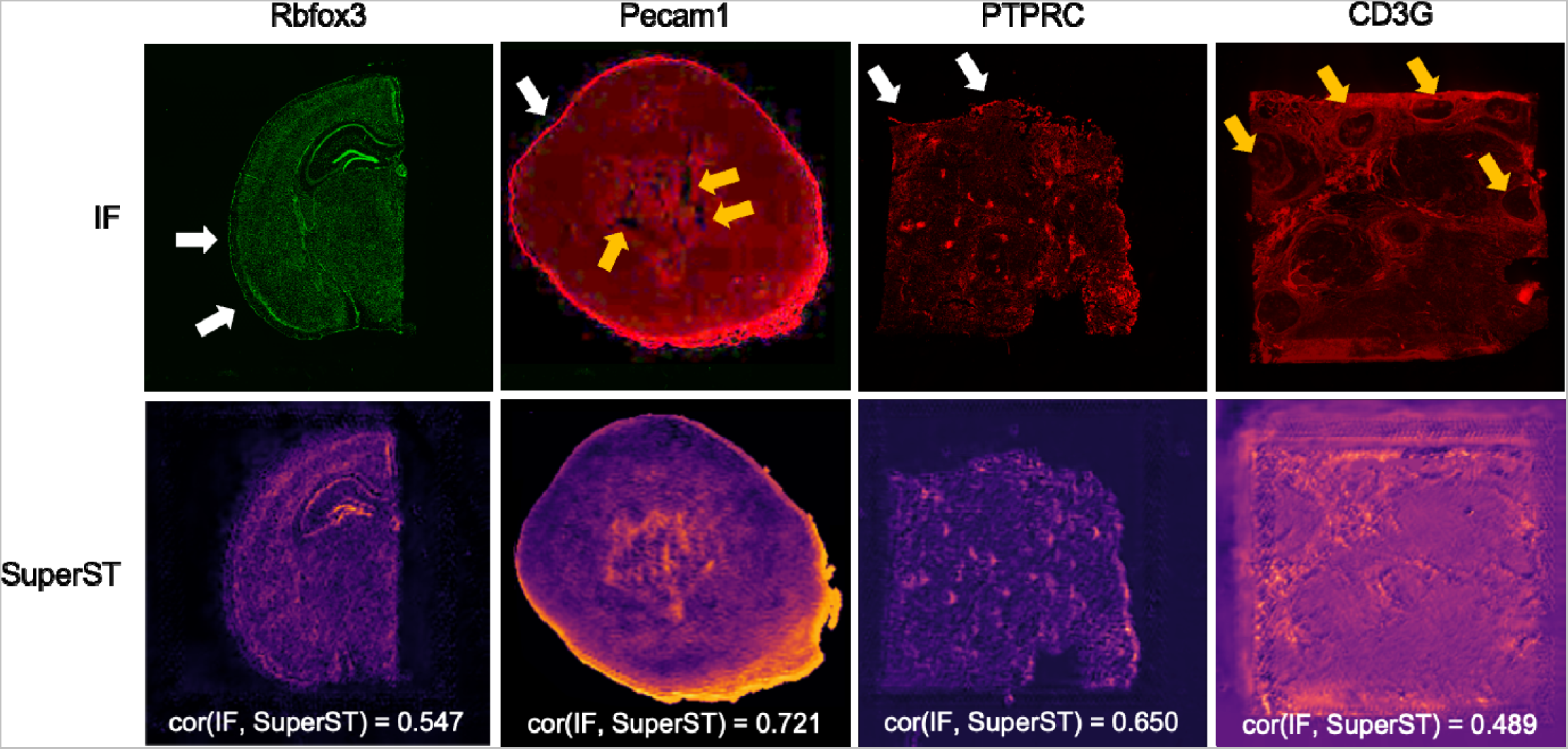

