## Supplementary Figures 1-4 for "Generation of Super-resolution Images from Barcode-based Spatial Transcriptomics Using Deep Image Prior"

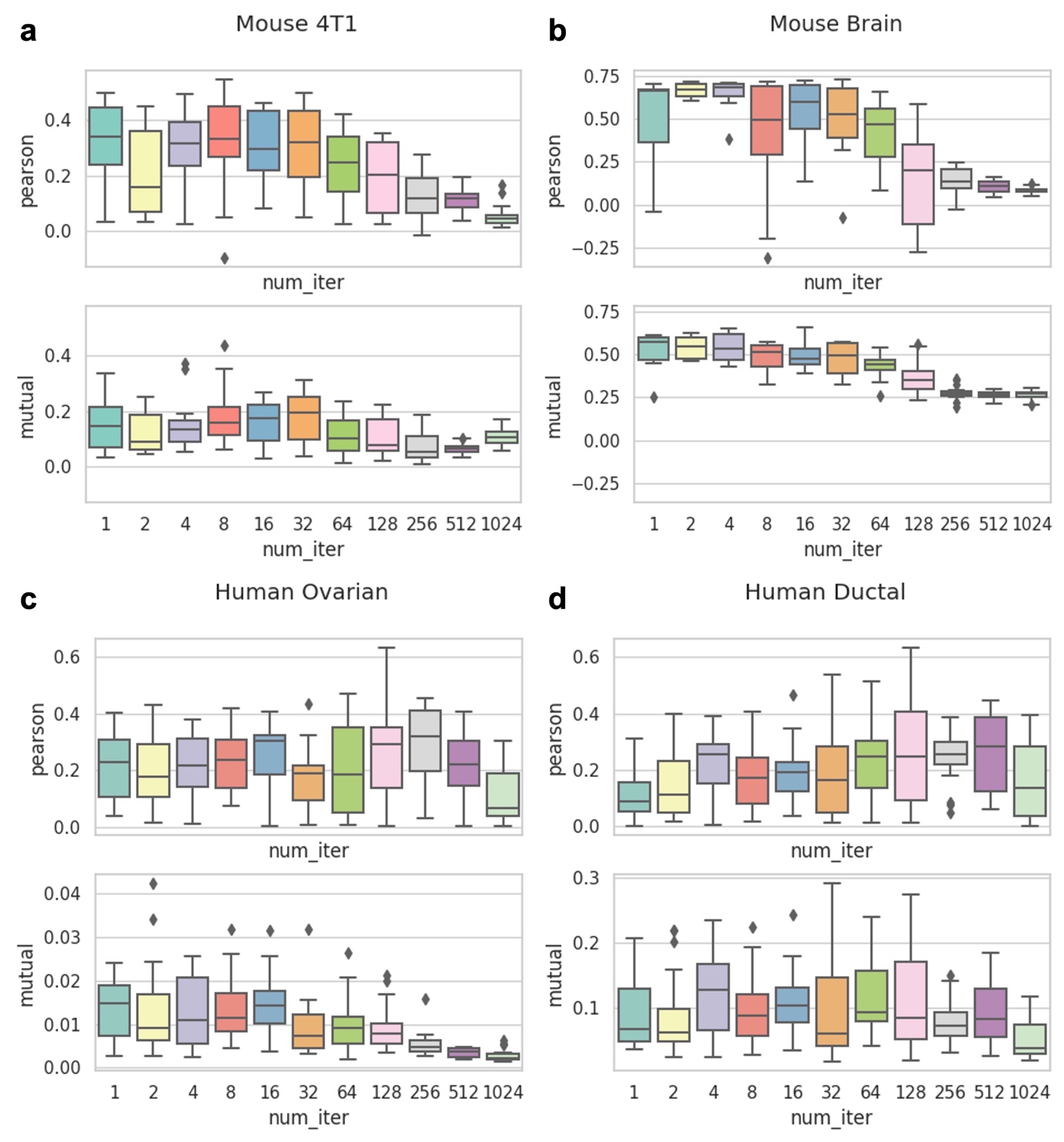

**Supplementary Figure 1** | Quantitative comparison between SuperST and IF. Image similarity metrics including Pearson correlation coefficient and mutual information were explored according to ‘*num_iter*’ in ‘*Mouse Brain*’ (**a**), ‘*Mouse 4T1*’ (**b**), ‘*Human Ovarian*’ (**c**), and ‘*Human Ductal*’ (**d**).

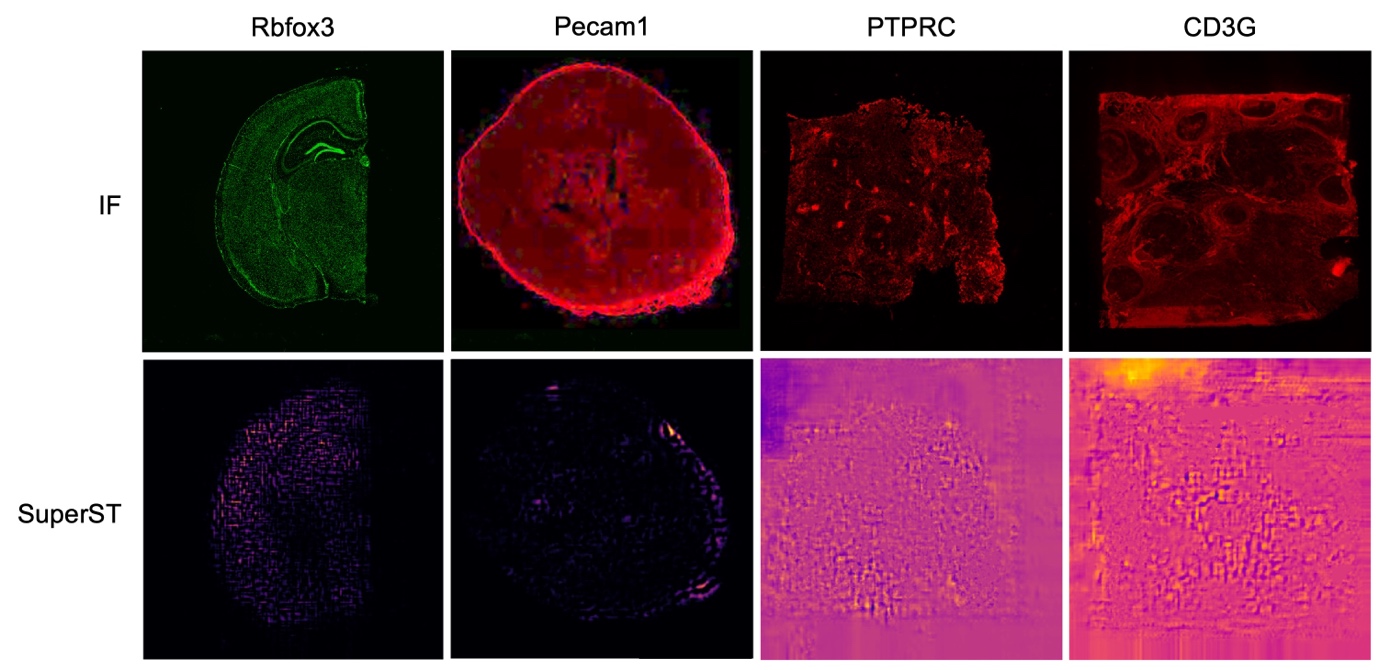

**Supplementary Figure 2** | The output images of SuperST for ‘*Mouse Brain*’, ‘*Mouse 4T1*’, ‘*Human Ovarian*’, and ‘*Human Ductal*’ when *‘num_iter*’ is 1024, along with the IF images. The model reacts excessively to sparse gene expression when the '*num_iter*' value exceeds 1024.

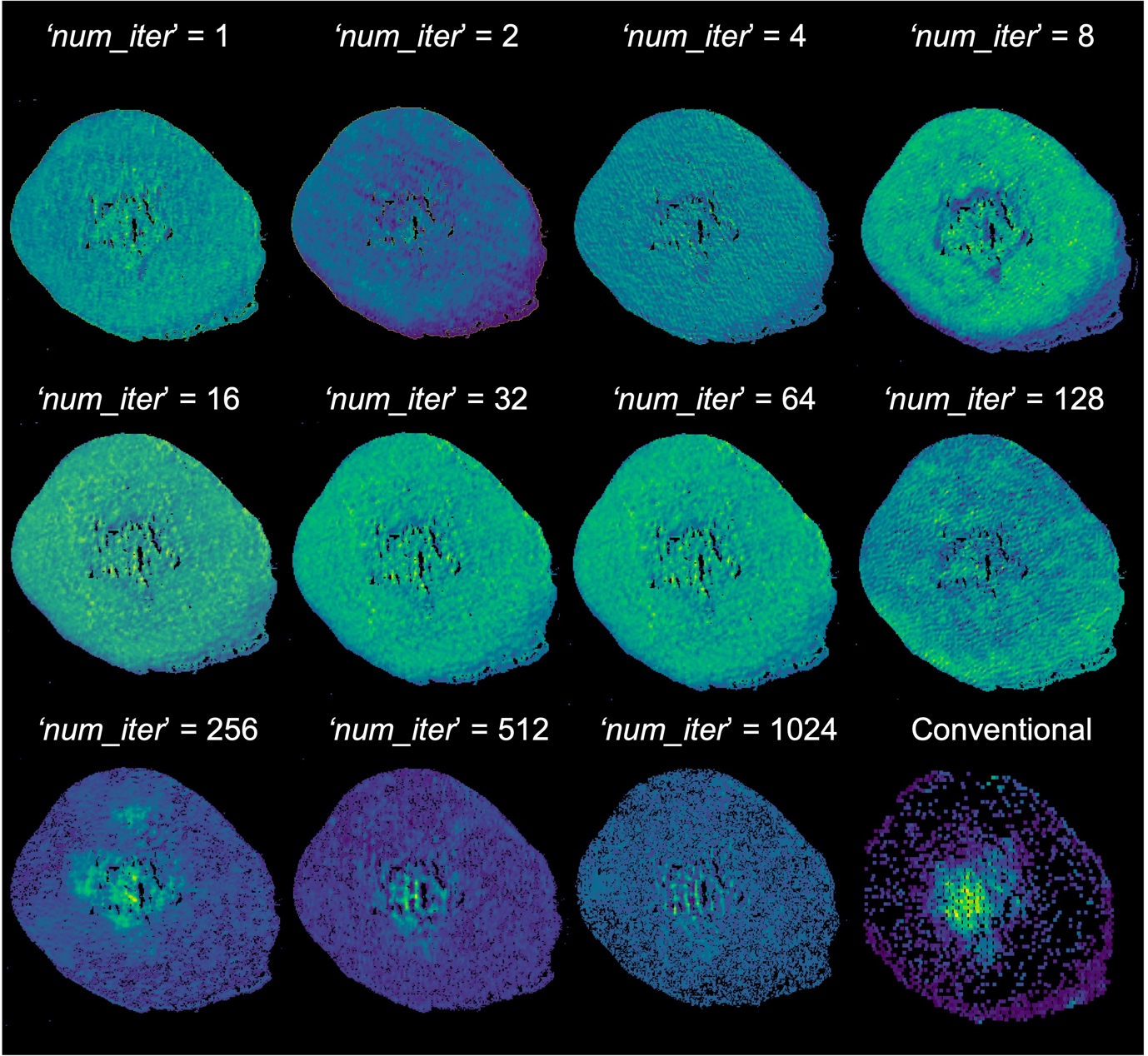

**Supplementary Figure 3** | The output images of SuperST for *S100a9* gene expression in ‘*Mouse 4T1*’ according to ‘*num_iter*’. We found that the output images displaying *S100a9* expression in the '*Mouse 4T1*' dataset achieved optimal performance when the '*num_iter*' parameter was set to 256.

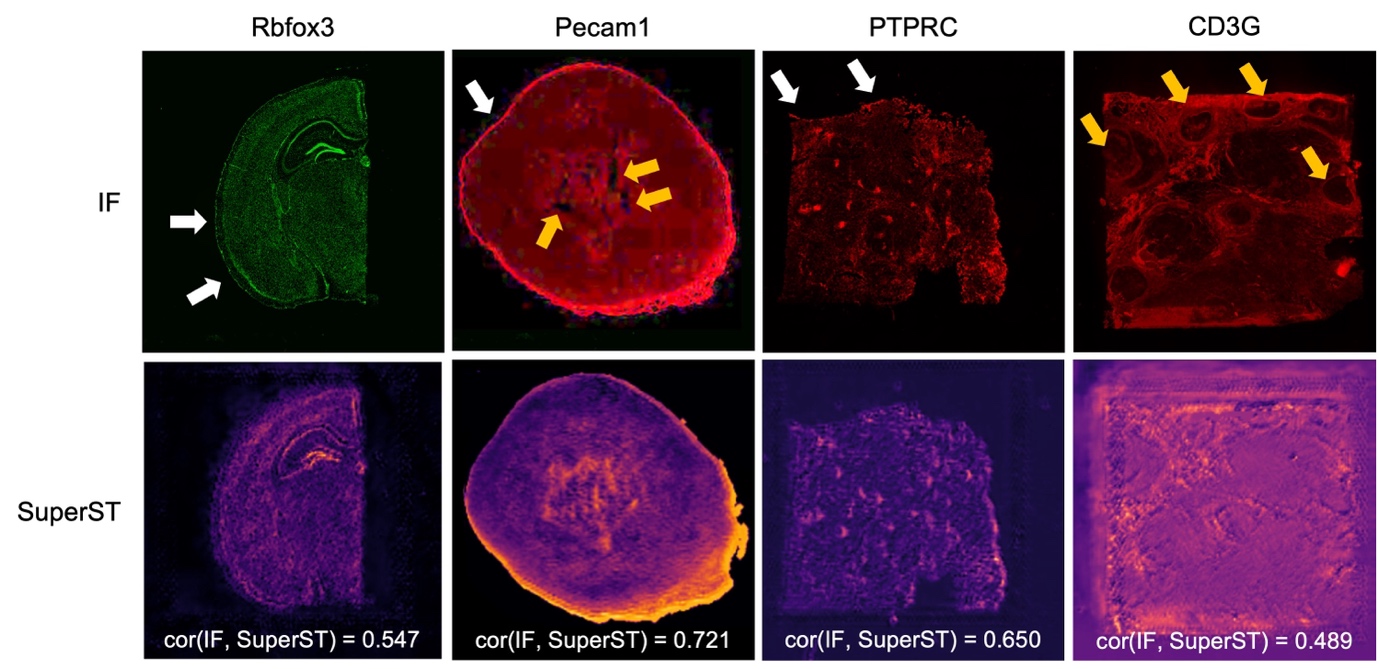

**Supplementary Figure 4** | Qualitative comparison between IF and SuperST. The samples of ‘*Mouse Brain*’, ‘*Mouse 4T1*’, ‘*Human Ovarian*’, and ‘*Human Ductal*’ were represented. White arrows showed the representative examples of autofluorescence patterns at tissue boundaries and yellow ones exhibited the effect of the histological conditions (*e.g.,* scarred tissue and duct-specific patterns) in IF images.
